# An integrative omics approach reveals posttranscriptional mechanisms underlying circadian temperature compensation

**DOI:** 10.1101/2021.10.06.463236

**Authors:** Christoph Schmal, Bert Maier, Reut Ashwal-Fluss, Osnat Bartok, Anna-Marie Finger, Tanja Bange, Stella Koutsouli, Maria S. Robles, Sebastian Kadener, Hanspeter Herzel, Achim Kramer

## Abstract

A defining property of circadian clocks is temperature compensation, characterized by the resilience of circadian free-running periods against changes in environmental temperature. As an underlying mechanism, the balance or critical reaction hypothesis have been proposed. While the former supposes a temperature-dependent balancing of reactions with opposite effects on circadian period, the latter assumes an insensitivity of certain critical period determining regulations upon temperature changes. Posttranscriptional regulations such as temperature-sensitive alternative splicing or phosphorylation have been described as underlying reactions.

Here, we show that knockdown of cleavage and polyadenylation specificity factor subunit 6 (*CPSF6*), a key regulator of 3’-end cleavage and polyadenylation, abolishes circadian temperature compensation in U-2 OS cells. We apply a combination of 3’-End-RNA-seq and mass spectrometry-based proteomics to globally quantify changes in 3’ UTR length as well as gene and protein expression between wild type and *CPSF6* knock-down cells and their dependency on temperature. Analyzing differential responses upon temperature changes in wild type and *CPSF6* knockdown cells reveals candidate genes underlying circadian temperature compensation. We identify that eukaryotic translation initiation factor 2 subunit 1 (*EIF2S1*) is among these candidates. *EIF2S1* is known as a master regulator of cellular stress responses that additionally regulates circadian rhythms. We show that knockdown of *EIF2S1* furthermore impairs temperature compensation, suggesting that the role of *CPSF6* in temperature compensation may be mediated by its regulation of *EIF2S1*.

## 1 Introduction

Circadian clocks are heritable endogenous oscillators that have evolved convergently across different taxa of life as adaptions to environments with predictable daily changes, such as light intensity and temperature (Bünning (1932); Rosbash (2009); Nikhil et al. (2020)). A defining feature of circadian clocks is that their rhythmicity is self-sustained, even under constant environmental conditions, with free-running periods of approximately (but not exactly) 24 hours. Remarkably, this rhythmicity is generated cell-autonomously by molecular delayed interconnected negative feedback loops (Bell-Pedersen et al. (2005)). In mammals, the core loop relies on the rhythmic transcriptional activation of *Per* and *Cry* genes by CLOCK and BMAL1 heterodimers, followed by the antagonistic effect of PER and CRY proteins on their own transcription (Ko and Takahashi (2006)). The interlocking of the core loop with additional positive and negative feedback loops is believed to give the system both plasticity and robustness (Akman et al. (2010); Yan et al. (2014); Pett et al. (2018); Schmal et al. (2019)). While the role of posttranslational modifications of clock proteins is considered crucial for circadian dynamics (Hirano et al. (2016)), there is emerging evidence that also posttranscriptional pre-mRNA processing steps such as capping, splicing, polyadenylation or nucleus-cytosol shuttling contributes to circadian gene expression (Preußner and Heyd (2016)).

Under natural conditions, environmental influences such as temperature, light intensity, or food availability can vary greatly with the time of day, but also at higher latitudes with season, and are subject to unpredictable fluctuations due to weather effects. Nevertheless, a functional clock system should reliably tell time even under such circumstances. Interestingly, although the rates at which chemical reactions occur generally increase substantially with temperature (Arrhenius (1889); Eyring (1935)), circadian free-running periods have been found to be remarkably resilient to changes in ambient temperature, typically exhibiting Q_10_ temperature coefficients between 0.85 to 1.2 (Sweeney and Hastings (1960)). This phenomenon, termed temperature compensation, is a defining feature of circadian clocks (Pittendrigh (1960)) that has been found in various kingdoms of life, ranging from poikilothermic organisms such as single-cell algae (Bruce and Pittendrigh (1956); Hastings and Sweeney (1957)), higher plants (Gould et al. (2006)), fungi (Sargent et al. (1966); Gardner and Feldman (1981)), insects (Pittendrigh (1954)) and reptiles (Menaker and Wisner (1983)) to cells and tissues of homeothermic animals such as rodents (Ruby et al. (1999); Izumo et al. (2003)) and human (O’Neill and Reddy (2011)). In 1973, Pittendrigh and Caldarola proposed that temperature compensation is only one aspect of an overarching homeostatic mechanism that protects the dynamics of the circadian clock from external influences (Pittendrigh and Caldarola (1973)) including metabolic fluctuations (Sancar et al. (2012); Johnson and Egli (2014)). In addition, temperature compensation among other pacemakers such as lunar and semi-lunar clocks (Kaiser and Neumann (2021)) has been reported, underpinning its importance for reliable time-telling even across different time scales.

Although temperature compensation of circadian rhythms was first described nearly a century ago, the underlying molecular mechanisms and regulatory elements involved remain poorly understood. Two conceptually different hypotheses for temperature compensation mechanisms have been proposed. (i) The balance hypothesis, i.e., counterbalance of the period responses of two or more temperature-dependent reactions leads to period stability (Hastings and Sweeney (1957); Pavlidis et al. (1968); Lakin-Thomas et al. (1991); Ruoff (1992); Domijan and Rand (2011); Kurosawa et al. (2017)). (ii) The critical reaction hypothesis, i.e., temperature insensitivity of certain critical period-determining processes leads to period insensitivity to temperature (Hong and Tyson (1997); Hong et al. (2007); Isojima et al. (2009); Shinohara et al. (2017); Narasimamurthy and Virshup (2017)). Interestingly, it has been reported that posttranscriptional mechanisms such as thermosensitive promoter use, alternative splicing and control of translation rates mediate both temperature responses and temperature compensation (Liu et al. (1997); Majercak et al. (1999); Mehra et al. (2009); Portolés and Más (2010); James et al. (2012); Narasimamurthy and Virshup (2017); Martin Anduaga et al. (2019); Chung et al. (2020)).

Here, we identified thermosensitive use of alternative polyadenylation sites as a possible mechanism underlying circadian temperature compensation in the human circadian clock. Using a systematic genetic screen, we found that knockdown of cleavage and polyadenylation specificity factor subunit 6 (*CPSF6*) leads to both long circadian periods and impaired circadian temperature compensation in human U-2 OS cells. *CPSF6* is a core component of the cleavage and polyadenylation machinery required for the utilization of distal polyadenylation sites and thus for normal lengths of 3’ UTRs. Using a global integrative multi-omics analysis comparing the differential temperature response in wild-type and *CPSF6* knockdown cells at the level of 3’ UTR length, transcript and protein expression, we identified candidate genes at the core of circadian temperature compensation.

## 2 Results

### 2.1 CPSF6 regulates circadian temperature compensation

To search for genes that regulate circadian temperature compensation via posttranscriptional mechanisms, we performed a large-scale RNA interference screen in human U-2 OS cells, an established model of the mammalian circadian clock (Maier et al. (2009, 2021)). Regardless of whether the balance hypothesis or the critical reaction hypothesis holds, depletion of essential components underlying temperature compensation should lead to a circadian period phenotype at least at certain temperatures. Therefore, we depleted the transcripts of 1.024 genes associated with posttranscriptional regulation and RNA processing (Carbon et al. (2008)) in U-2 OS reporter cells with RNAi constructs from our laboratory library and analyzed the circadian bioluminescence rhythms at 35°C. Knockdown of 83 genes showed significant effects on circadian periods (p < 0.01; Tab. S1). Depletion of *CPSF6* resulted in the strongest phenotype, i.e. a dose-dependent (Fig. S1) period lengthening of approximately 1.5-2 hours (Fig. 1A) in both human U-2 OS and mouse NIH3T3 cells (Fig. S1). *CPSF6* is part of a multi-protein complex that is important for 3’-end processing of transcripts defining the polyadenylation site and thus the 3’ UTR length. Interestingly, knockdown of NUDT21 (also known as CPSF5), a heterodimeric binding partner of CPSF6 phenocopies the effect of *CPSF6* (Fig. S1), suggesting that *CPSF6* acts on the circadian clock via its role within the polyadenylation complex.

**Figure 1:**
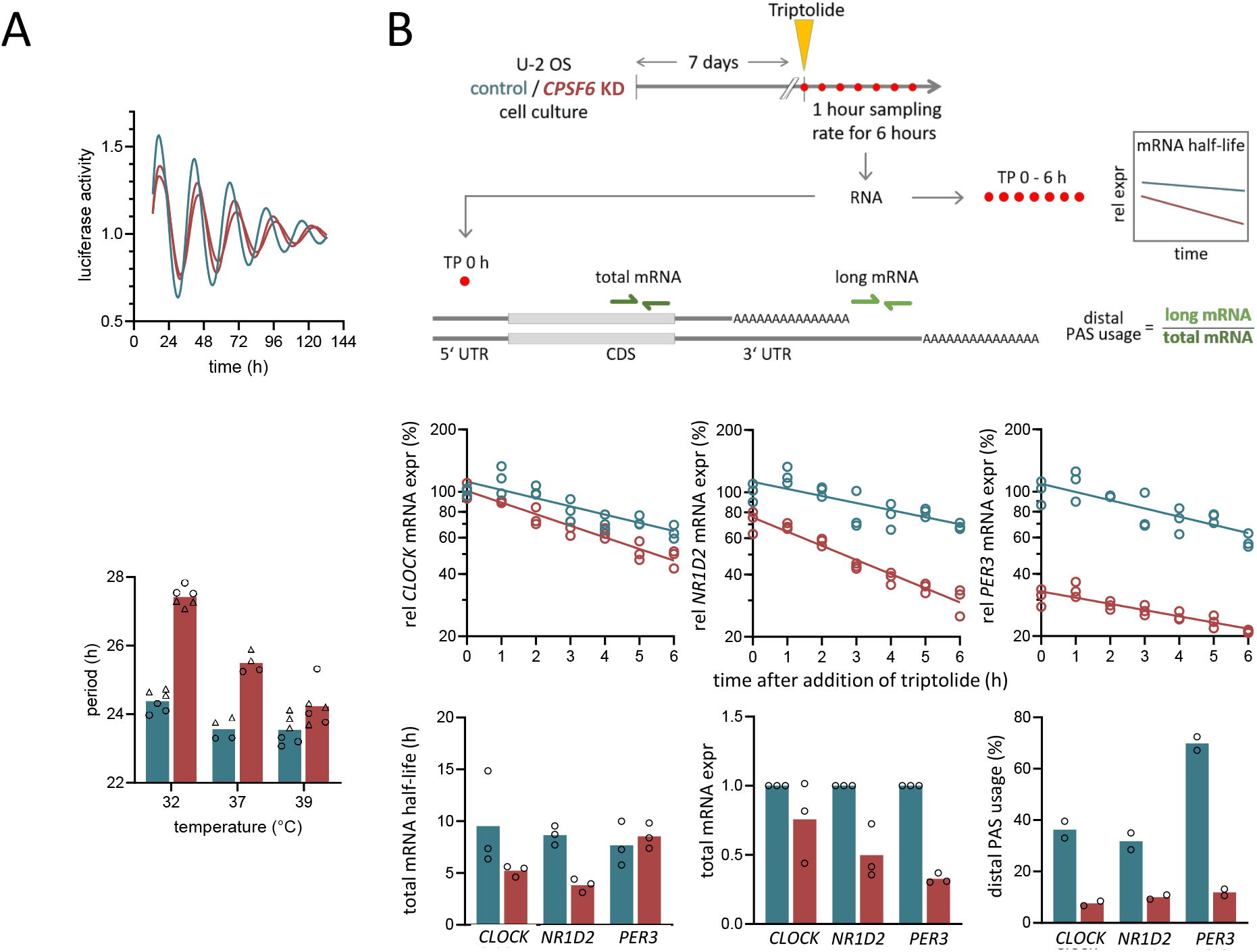
Knockdown of *CPSF6* impairs circadian temperature compensation. A) Circadian free-running period of U-2 OS cells lengthens upon RNAi-mediated *Cpsf6* knockdown (*red*) in comparison with control cells (*blue*), as accessed by a *Bmal1-luciferase* reporter construct. Raw time series data has been normalized by its baseline trend (magnitude). B) *Upper panel:* Schematic drawing of experimental setup for determination of alternative polyadenylation site usage, transcript expression as well as transcript half-lives. *Middle panel*: Dynamics of *Clock* (*left*), *Nr1d2* (*middle*) and *Per3* (*right*) mRNA expression after application of *triptolide* in wild type (*blue*) and *CPSF6* knockdown cells (*red*). Straight lines depict linear fits to the logarithmic data as used for the determination of half-lives. *Bottom panel*: Bar plots, comparing the total mRNA half-life (*left*), total mRNA expression (*middle*) and distal polyadenylation site (PAS) usage (*right*) in *Clock, Nr1d2,* and *Per3* in wild type versus *CPSF6* knockdown cells. Dots denote raw data while bars denote corresponding averages. C) Temperature compensation is significantly reduced upon *Cpsf6* knockdown.

Since 3’ UTRs contribute to the rhythmic expression of clock and clock-controlled genes (Kwak et al. (2006); Woo et al. (2009)) and thermosensitive regulation of pre-mRNA processing has been proposed as a mechanism underlying temperature compensation in other organisms, we investigated the effect of temperature on free-running periods in *CPSF6* knockdown cells. Interestingly, knockdown of *CPSF6* significantly affects temperature compensation. Whereas the free-running periods of control cells changed only slightly from 24.4h ± 0.3h at 32°C to 23.5h ± 0.4h at 39°C, the temperature dependence was significantly altered upon *CPSF6* knockdown with a shortening of free-running period from 27.4h ± 0.3h at 32°C to 24.2h ± 0.6h at 39°C (Fig. 1C, Fig. S3A). These dependencies led to an increase of the period temperature coefficient from 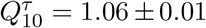 to 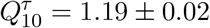 upon *CPSF6* knockdown (Fig. S3B).

#### CPSF6 knockdown alters mRNA stability and expression of clock genes

It has been described that *CPSF6* knockdown induces a systematic shift toward proximal polyadenylation sites (PAS) and thus to shorter 3’ UTRs (Martin et al. (2012)). To investigate, whether *CPSF6* knockdown has an impact on 3’ UTR length of canonical clock genes, we measured the expression of long 3’ UTR isoforms versus the total amount of mRNA. Among the nine clock genes examined, *ARNTL*, *CRY1/2*, *CLOCK*, *NR1D1/2* and *PER1/2/3*, transcripts of *PER3* showed the largest shift from distal to proximal PAS usage at 37°C, followed by *CLOCK* and *NR1D2* (Fig. 1B). Whereas *CRY2*, *NR1D1* as well as *PER1* showed no significant change in polyadenylation-site usage, *CPSF6* knockdown resulted in a small increase of distal polyadenylation-site usage for *CRY1* and *PER2* (Fig. S2).

Since 3’ UTRs have been shown to affect mRNA stability (Mayr (2017)), we compared the mRNA half-lives of clock genes in unsynchronized control and *CPSF6* knockdown cells at 37°C (Fig. 1B) (Bensaude (2011)). For *CLOCK* and *NR1D2*, which showed a significant shift toward shorter 3’ UTRs, we observed reduced mRNA half-lives (Fig. 1B), in contrast to the expectation that a lack of miRNA binding sites in transcripts with short 3’ UTRs generally leads to an increased stability (Sandberg et al. (2008)). However, *PER3*, which also had an overall shorter 3’ UTR in *CPSF6* knockdown cells, showed no changes in mRNA stability (Fig. 1B). In all three cases, *CPSF6* knockdown reduced total mRNA expression. Thus, whereas shortening of the 3’ UTR length in *CLOCK* and *NR1D2* results in shorter transcript half-lives, presumably through a loss of regulatory elements within the UTR, our data suggest that processes other than mRNA stability regulate changes of *PER3* expression.

Of note, we did not detect a systematic temperature effect on the 3’ UTR lengths of clock genes, nor was the *CPSF6* knockdown effect on the 3’ UTR lengths of the clock genes examined temperature dependent (not shown). This suggests that the role of *CPSF6* in circadian temperature compensation is not limited to the core oscillator, but may be a higher-level mechanism.

### 2.2 *CPSF6* knockdown induces global changes in polyadenylation site usage, transcript and protein expression

The increased temperature sensitivity of free-running periods among *CPSF6* knockdown cells suggests regulatory interactions at the pre-mRNA processing level as a potential mechanism underlying circadian temperature compensation in mammals. By combining transcriptomics, proteomics as well as RNAi high-throughput approaches, we aim to uncover key regulatory components that contribute to the observed differential temperature response between *CPSF6* knockdown and wild type cells and thus temperature compensation. To investigate the impact of temperature and *CPSF6* on polyadenylation site selection, total transcript and protein expression, we employed an integrative omics approach in *CPSF6* knockdown and control cells at three different temperatures (32°C, 37°C, and 39°C) (Fig. 2A). Below, we first describe the effect of *CPSF6* knockdown on polyadenylation site usage, transcript and protein abundance at one temperature (37°C), before reporting whether and to what extent temperature modulates these effects.

**Figure 2:**
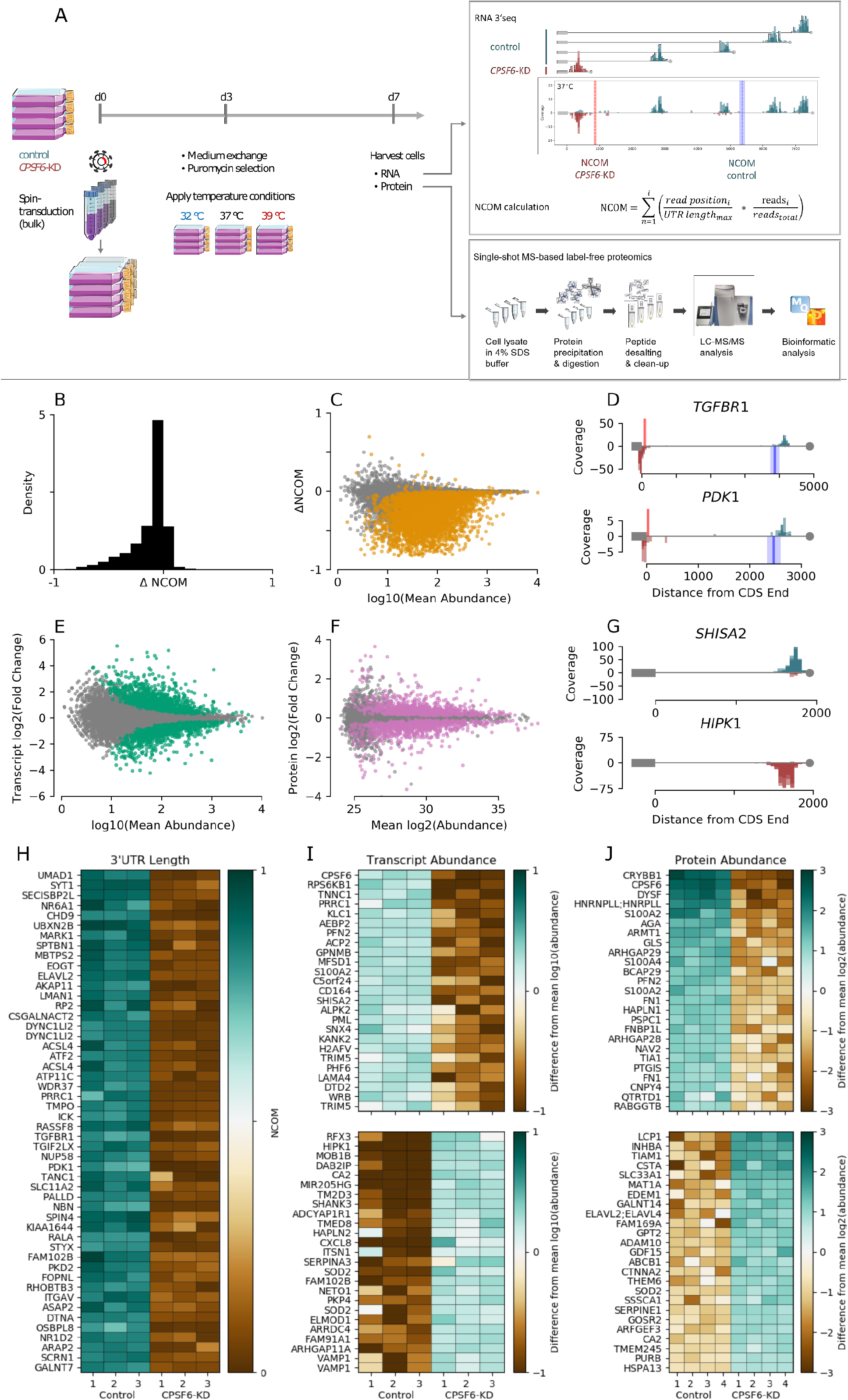
Knockdown of *CPSF6* leads to global shifts in polyadenylation site usage as well as major transcript and protein abundance changes. A) Schematic of the integrative omics experimental approach used throughout this study. B) Histogram depicting the absolute shift in NCOM (i.e. normalized 3’ UTR length) upon *CPSF6* knockdown at 37°C. C) Absolute shift in NCOM upon *CPSF6* knockdown versus decadic logarithm of the corresponding mean expression across both genetic backgrounds. Isoforms showing a significant NCOM shift (t-test) are depicted by *orange dots* (Benjamini-Hochberg corrected *p* < 0.05). D) Two exemplary 3’ UTR read distributions of genes showing a significant shortening of 3’ UTR length upon *CPSF6* knockdown (*red*) in comparison to wild type (*blue*). Note that the three replicates are overdubbed. Bold vertical lines and attached shaded areas, denote the NCOM average and standard deviation among the replicates, respectively. E) MA plot showing the binary logarithm of the fold change between wild type and *CPSF6* knockdown transcript expression versus the decadic logarithm of the corresponding mean expression. Statistically significant differential expressions as determined via DESeq are denoted by green markers (Benjamini-Hochberg corrected *FDR* < 0.05). F) MA plot showing the binary logarithm of fold changes between wild type and *CPSF6* knockdown cells versus corresponding mean protein LFQ intensities, which we will thereinafter term protein abundances for the sake of simplicity. Proteins showing a significantly different abundance (t-test) are depicted by *pink dots* (Benjamini-Hochberg corrected p < 0.05). G) Representative examples that show a decrease (SHISA2, *top*) or increase (HIPK1, *bottom*) of transcript abundance upon *CPSF6* knockdown. H) Heatmap depicting NCOM values for the 50 isoforms with the largest shift among those transcripts that show a significant NCOM change upon *CPSF6* knockdown. I) Heatmap showing 25 transcript isoforms with a significant negative (top) and positive (*bottom*) fold change upon *CPSF6* knockdown, sorted by the amount of fold change. J) Same as panel (I) for protein LFQ abundance.

#### Polyadenylation site usage is globally controlled by CPSF6

As a prerequisite for further analyses, we first precisely mapped polyadenylation sites of transcripts in U-2 OS cells by means of an RNaseH-seq approach as previously described (Afik et al. (2017)). Of all ≈20.000 genes for which we found polyadenylation signals, the vast majority (> 70%) contained multiple polyadenylation signals (Fig. S4, Tab. S2), similar to what has been reported for other tissues (Derti et al. (2012)). To investigate the impact of *CPSF6* on polyadenylation site usage, we performed 3’-end RNA-seq and calculated the ‘‘center of mass” of the read distribution along the annotated 3’ UTR, normalized to the lengths of the 3’ UTR. Hereafter, we refer to this as the normalized center of mass (NCOM, see Fig. 2A). Knockdown of *CPSF6* resulted in a global shift toward use of proximal polyadenylation sites leading to overall shorter 3’ UTRs at 37°C (Fig. 2B, Tab. S3). A significant shift of the polyadenylation site was observed in approximately one-third of all ≈ 12.000 isoforms detected (Fig. 2C *orange dots* and Fig. 2H), with the majority (99%) expressing a short 3’ UTR transcript variant, which is very similar to previous reports (Martin et al. (2012); Gruber et al. (2012); Li et al. (2015)). Figure 2D illustrates two representative examples of read distributions of transcripts showing a significant switch from expression of long 3’ UTR isoforms in wild type (*blue*) to expression of short 3’ UTR isoforms in *CPSF6* knockdown cells (*red*).

#### Transcript and protein abundance are globally controlled by CPSF6

To study the functional consequences of alternative polyadenylation, we next quantified changes in gene expression after *CPSF6* knockdown. We detected 1.924 (16%) differentially expressed transcripts (Fig. 2E *green dots*; Fig. 2G shows two examples), of which 977 were upregulated and 947 were downregulated (Fig. 2I, Tab. S4). Interestingly, these transcripts were enriched within the group of transcripts that also exhibited a significant shift in 3’ UTR length (Fig. 3A *light green*), indicating a dominant role of 3’ UTR length in gene expression, as previously reported (Mayr (2017)). Out of the 967 transcripts that differed significantly at both levels, 60% showed an increase and 40% a decrease in gene expression, respectively. This is consistent with recent findings reporting that a reduction in 3’ UTR length can lead to both increased or decreased gene expression (Mayr (2017)), in contrast to the previously prevailing assumption that a shorter 3’ UTR generally leads to an increased mRNA stability by loosing miRNA binding sites.

**Figure 3:**
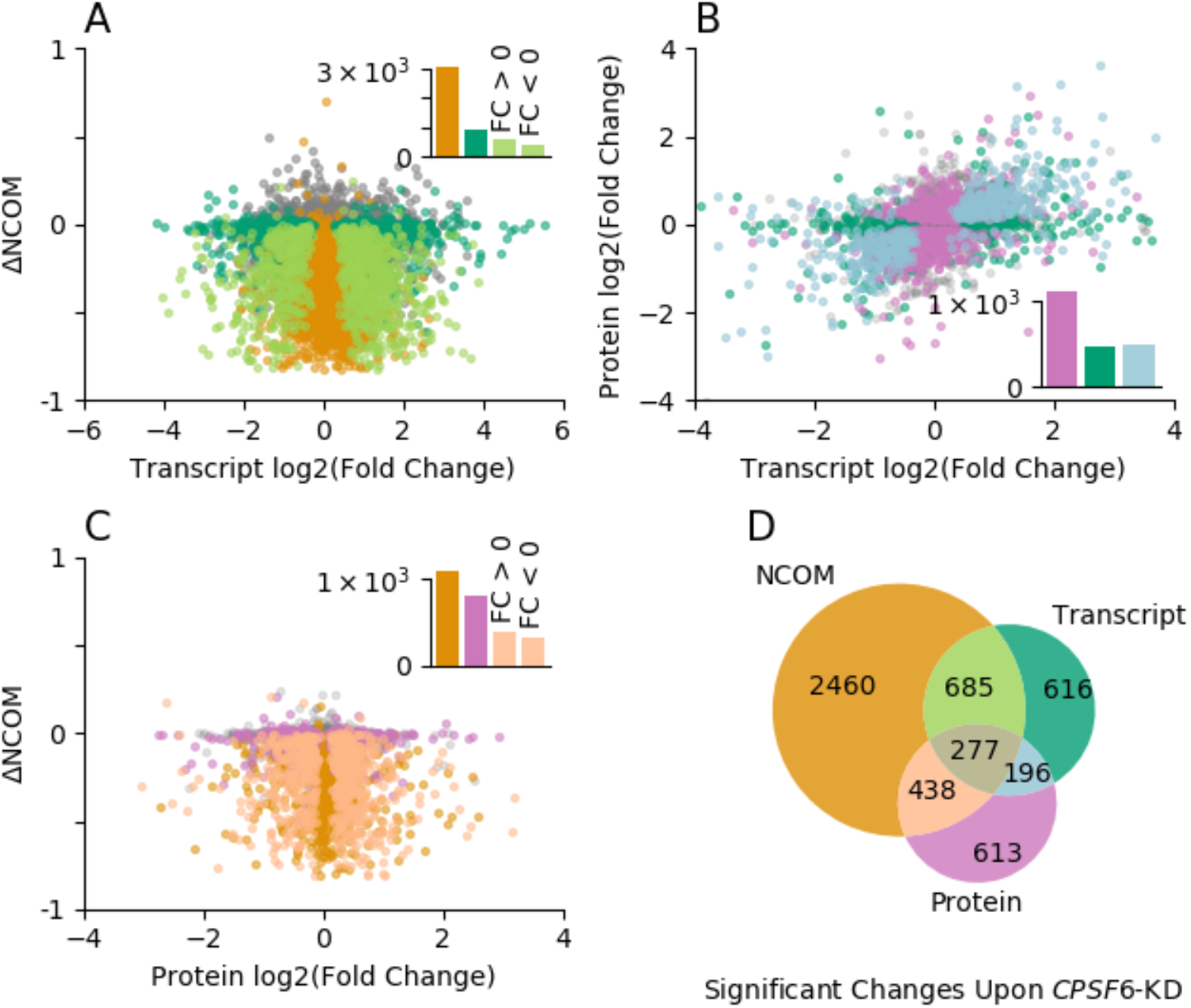
Global shortening of 3’ UTR lengths upon *CPSF6* knockdown induces global changes in transcript and protein abundance. A) Scatter plot depicting the shift in 3’ UTR lengths (ΔNCOM) versus binary logarithm of fold changes in transcript expression upon *CPSF6* knockdown at 37° C. *ΔNCOM* is defined as the *normalized center of mass* (NCOM) in wild type U-2 OS cells, subtracted from the NCOM in *CPSF6* knockdown cells. Different colors denote transcripts with no significant changes in NCOM and transcript abundance (*gray*), significant changes exclusively at the NCOM (*orange*) or transcript (*dark green*) level or at both levels at the same time (*light green*), respectively. Transcripts with a significant fold change upon *CPSF6* knockdown are enriched within the set of transcripts that show a significant shift in 3’ UTR length (Fisher’s exact test: *p* < 0.001, odds-ratio 2.24). B) Binary logarithm of protein fold changes upon *CPSF6* knockdown are depicted versus corresponding binary logarithms of fold changes of transcript expression. Different colors denote genes with no significant changes in transcript and protein expression (*gray*), solely in transcript abundance (*dark green*), solely in protein abundance (*pink*) or in transcript and protein abundance (*light blue*), respectively. Genes with a significant fold change at the protein level are significantly enriched within the set of genes that exhibit a significant fold change of transcript expression (Fisher’s exact test: p < 0.001, odds-ratio 2.38). C) Shift in 3’ UTR lengths (ΔNCOM) versus binary logarithm of fold change in protein abundance upon *CPSF6* knockdown. Different colors denote genes with no significant change in 3’ UTR length and protein abundance (*gray*), solely in 3’ UTR length (*orange*), solely in protein levels (*pink*) or a significant change in both (*light orange*), respectively. Proteins whose intensities are significantly different between conditions (fold change) are moderately enriched within the set of transcripts exhibiting a significant change in 3’ UTR length upon *CPSF6* knockdown (Fisher’s exact test: p = 0.0002, odds-ratio 1.27). D) Venn diagram indicating the number of genes that show a significant alteration of 3’ UTR length (*orange*), transcript expression (*green*) or protein abundance (*pink*) upon *CPSF6* knockdown. Note that significance thresholds identical to those in Fig. 2 have been used. *Insets* in panels A-C depict the absolute number of isoforms in each corresponding group.

To gain further insight into the functional implications of *CPSF6* knockdown, we globally examined how *CPSF6* dependent shortening of 3’ UTRs affects protein abundance (Fig. 2A). Using mass spectrometry-based quantitative proteomics, we quantified 4.733 proteins across all samples. Of these, 1.683 (36%) showed a significantly different abundance after *CPSF6* knockdown (Fig. 2F *pink*), with 50% upregulated and 50% downregulated (Fig. 2J, Tab. S5). These altered proteincoding genes were significantly enriched in the group of genes that also showed significant regulation at the transcriptional level (Fig. 3B), indicating a high correlation between mRNA and protein expression, as previously described (Schwanhäusser et al. (2011)). Although still significant, genes whose protein level was significantly altered upon *CPSF6* knockdown were enriched to a much lesser extent in the set of transcripts that exhibited a change in 3’ UTR length after *CPSF6* knockdown (Fig. 3C), indicating a relatively weaker effect of global 3’ UTR shortening on protein abundance, similar to what has been previously described for human and murine T cells (Gruber et al. (2014)).

In summary, knockdown of *CPSF6* induces global changes in 3’ UTR length as well as in transcript and protein abundance at physiological temperatures of 37°C.

### 2.3 Temperature has little effect on global alternative polyadenylation but major influence on transcript and protein abundance

To identify genes that control circadian temperature compensation in wild-type cells or analogously promote temperature decompensation in *CPSF6* knockdown cells, we next examined the extent to which temperature induces genome-wide expression changes in polyadenylation site usage as well as transcript and protein abundance. Interestingly, only minor alterations in polyadenylation site selection were observed upon changes in temperature in wild-type cells, especially compared to the substantial global shifts seen upon *CPSF6* knockdown (compare Fig. 2B, C and 4A). Of the ≈ 12.000 isoforms analyzed, only 2% showed a significant temperature-mediated change in 3’ UTR length (Fig. 4A *orange*, Fig. S6A, Tab. S6). Of these, > 75% showed an increase in NCOM with increasing temperatures, indicating that higher temperatures tend to elongate 3’ UTRs in U-2 OS cells. This is in contrast to previous reports that cold shock induces a switch towards longer 3’ UTRs in mouse embryonic fibroblasts (Liu et al. (2013)).

**Figure 4:**
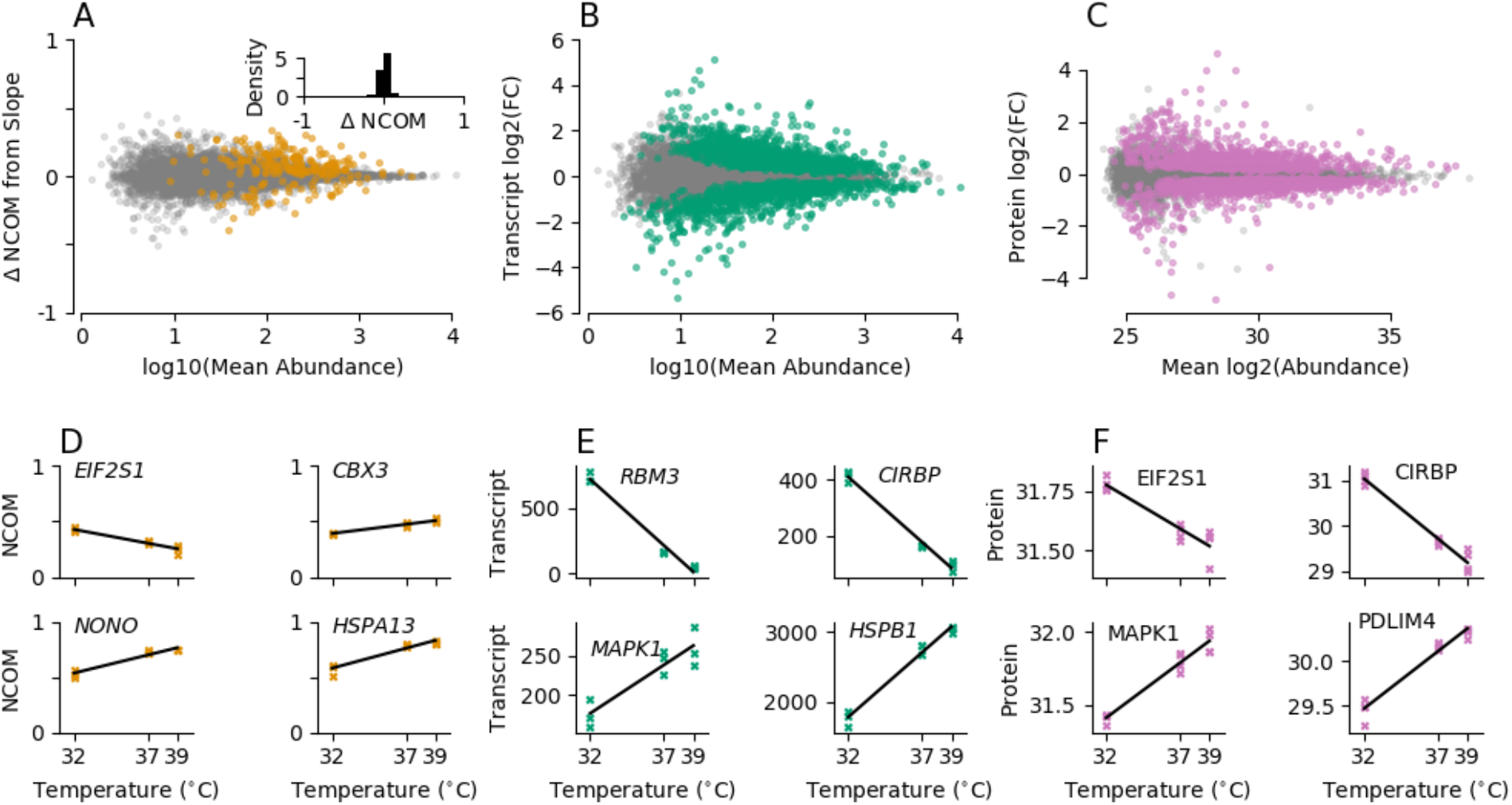
Alterations in environmental temperature induce major expression changes while having a weak effect on 3’ UTR length. A) MA plot showing the shift in 3’ UTR length upon temperature variations between 32°C and 39°C as determined from a linear regression versus the decadic logarithm of the mean expression among all temperatures from the corresponding transcript. Transcripts with a statistically significant change in 3’ UTR length at a 5% FDR (Benjamini-Hochberg corrected p-values) are depicted by *orange* markers. No global shift in 3’ UTR length can be observed upon temperature alterations in wild type cells (*inset*). B) Same as panel (A), showing the binary logarithm of transcript expression fold changes upon temperature variations between 32°C and 39°C on the ordinate. Transcripts with a statistically significant change in expression at a 5% FDR (Benjamini-Hochberg corrected p-values) are depicted by *green markers.* Same as panel (B), showing the binary logarithm of protein LFQ intensity fold changes upon temperature variations between 32°C and 39°C on the ordinate as well as the binary logarithm of the mean among all temperatures from the corresponding proteins. Proteins with a statistically significant change in abundance at a 5% FDR (Benjamini-Hochberg corrected p-values) are depicted by *pink markers*. D-F) 3’ UTR length (D, *orange markers*), transcript expression (E, *green markers*) and protein intensities (F, *pink markers*) in wild type cells at all three experimentally measured temperatures for different example genes that show a statistically significant corresponding temperature response. Fits from a linear regression are depicted by black lines.

Despite the relatively weak effect on 3’ UTR lengths, changes in ambient temperature resulted in major expression changes at the transcript and protein levels (Fig. 4B, C, Tab. S7, S8). A total of 2.445 (20%) transcripts showed a significantly different abundance upon temperature changes (Fig. 4B, Tab. S7). Of these, more than half (60%) showed an increased abundance at higher temperatures. Furthermore, 43% of the proteins show significant abundance changes across different temperatures with almost equal increases and decreases in abundance at higher temperatures (Fig. 4C, Tab. S8). A similar picture can be drawn for *CPSF6* knockdown cells in response to different ambient temperatures. As in wild-type cells, only a small fraction (1%) of transcripts had significantly different 3’ UTR lengths (Fig. S7A), whereas abundance changed substantially at the transcript (22%) and protein (50%) level (Fig. S7B,C).

Taken together, these data suggest that the cellular response to temperature is largely independent of *CPSF6*. However, we cannot exclude the possibility that temperature and *CPSF6* modulate 3’ UTR length and thus the expression of a master regulator of transcription and/or translation.

### 2.4 Toward identifying key players underlying temperature compensation

Regardless of whether the balance hypothesis or the critical reaction hypothesis holds, temperature decompensation in a particular genetic background (*e.g. CPSF6* knockdown) can be explained by a differential response to ambient temperatures compared to wild type behavior. Therefore, we searched for genes that showed a significant difference in the temperature-dependent changes of 3’ UTR length as well as transcript or protein abundance between wild-type and *CPSF6* knockdown cells. For simplicity, we have assumed a linear relationship between the observed variable of interest and temperature and analyzed whether such linear relationship was significantly different (Fig. S3A). We then converted the resulting slopes to Q_10_ temperature coefficients for comparison with previous results.

Comparing wild-type with *CPSF6* knockdown cells, we observed a differential response to temperature for 182 transcripts (2%, Fig. 5A *orange dots*, Tab. S9) in terms of 3’ UTR length and for 942 transcripts (10%, Fig. 5B *green dots*, Tab. S10) in terms of expression level. At the protein level, we detected 678 proteins (14%, Fig. 5C *pink dots*, Tab. S11) whose temperature-dependent abundance differed between wild-type and knockdown cells. Interestingly, only six genes (*CBX3*, also known as *HP1γ*, *EIF2S1*, *HSPA13*, *MAPK1*, also known as *ERK2*, *NAP1L1* and *PDLIM4*) showed a significantly differential temperature response between wild type and *CPSF6* knockdown cells at all three regulatory levels examined, i.e. 3’ UTR length, transcript abundance, and protein abundance (Fig. 5D-E and Fig. S8).

**Figure 5:**
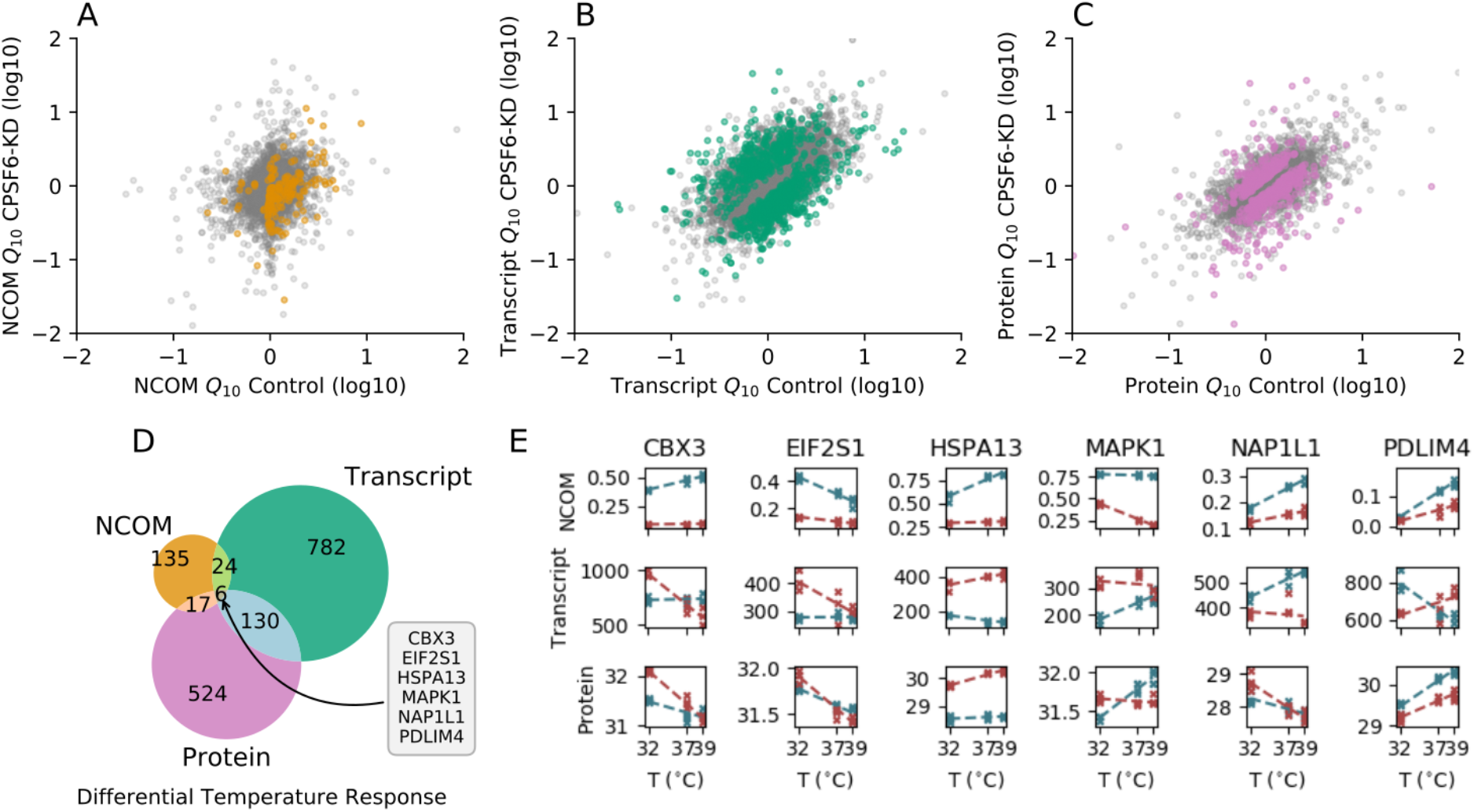
Differential temperature responses between wild type and *CPSF6* knock down cells suggests candidates underlying temperature compensation. A) NCOM temperature coefficient Q10 in *CPSF6* knockdown versus wild type cells. Significantly differential responses in polyadenylation site selection (i.e. NCOM shifts) upon temperature changes between wild type and *CPSF6* knockdown cells are depicted by *orange* dots. B) Transcript expression temperature coefficient Q_10_ in *CPSF6* knockdown versus wild type cells. Significantly different responses in transcript expression upon temperature changes between wild type and *CPSF6* knockdown cells are depicted by *green* dots. C) Protein abundance temperature coefficient Q_10_ in *CPSF6* knockdown versus wild type cells. Significantly different responses in protein expression upon temperature changes between wild type and *CPSF6* knockdown cells are depicted by *pink* dots. D) Venn diagram indicating the number of genes that show a statistically significant differential temperature response between wild type and *CPSF6* knockdown cells. E) 3’ UTR length (top *panels*), transcript expression (*middle panels*) as well as protein abundance (*lower panels*) for three different temperatures in wild type (*blue*) and *CPSF6* knockdown cells (*red*). Linear fits are denoted by dashed lines. Depicted are all six genes from panel (D) that show a differential temperature response between wild type and *CPSF6* knockdown cells at all three regulatory layers.

If any of these genes contribute to circadian temperature compensation, their genetic depletion should not only result in a circadian period phenotype, but this phenotype should also depend on temperature. Therefore, we first examined, which genes identified by our differential response analysis also showed a period lengthening or shortening phenotype after RNAi-mediated knockdown. Of the six genes that exhibited a differential temperature response at all three regulatory levels, only *EIF2S1* (Fig. 6A) and, to a much weaker extent, *HSPA13* (Fig. S9) showed a significant period change after RNAi-mediated knockdown. To test whether knockdown of *EIF2S1* also leads to impaired temperature compensation, similar to what has been observed with *CPSF6* knockdown (Fig. 1C), we compared the circadian periods at three different temperatures (Fig. 6B). Inter-estingly, knockdown of *EIF2S1* also resulted in temperature-decompensation and increased the period temperature coefficient from 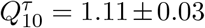 to 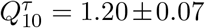 (Fig. 6C). This suggests that the role of *CPSF6* in circadian temperature compensation may be mediated via regulation of *EIF2S1*. Along these lines, a temperature dependent differential expression of isoforms with different 3’ UTR lengths may be responsible for buffering the corresponding transcript and protein abundance against temperature changes. In line with this assumption, predominant temperature insensitive expression of the short 3’ UTR isoform in *CPSF6* knockdown cells leads to a stronger temperature dependency of *EIF2S1* transcript and protein abundance (5 E, *second column*).

**Figure 6:**
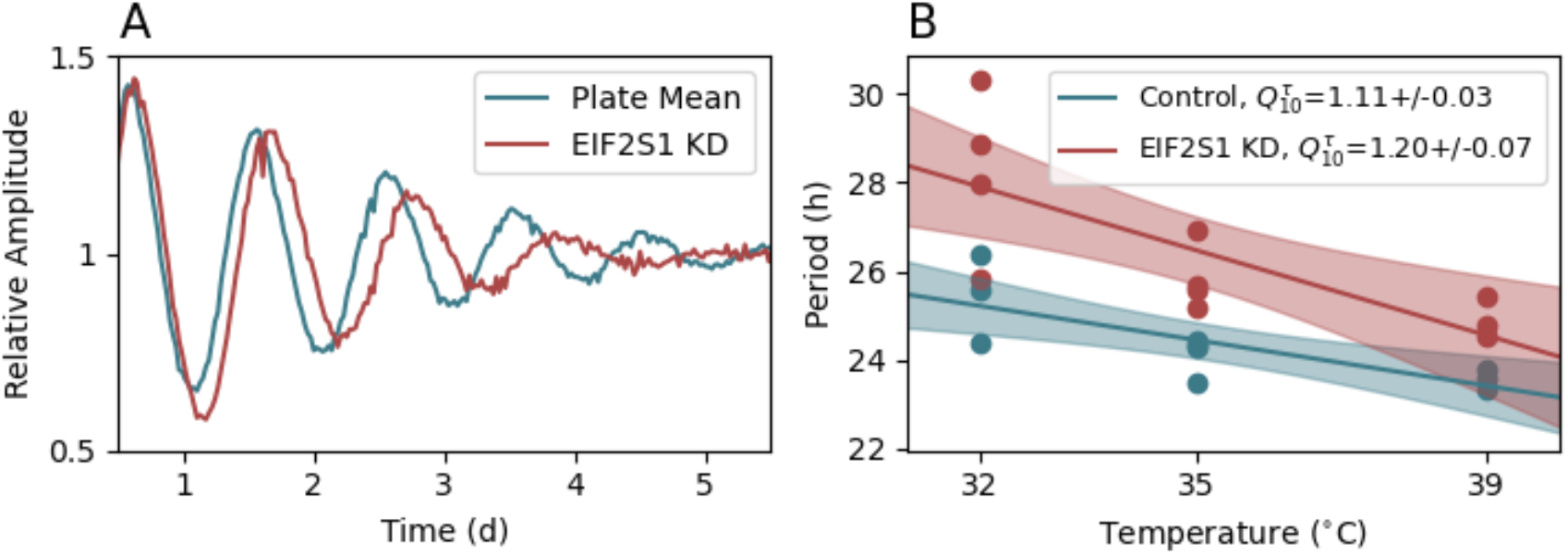
*EIF2S1* knockdown induces a long period phenotype and diminishes temperature compensation. A) Circadian phenotype of RNAi mediated knockdown of *EIF2S1* (*red*) together with the corresponding plate mean control (*blue*) of the corresponding large scale RNAi screen. B) Oscillatory periods of wild type (*blue*) and *EIF2S1* knockdown cells (*red*) at three different temperatures. Straight lines depict linear regressions as used to determined the corresponding period coefficient 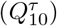.

## Discussion

In all organisms with circadian clocks, the period of circadian oscillations is resilient to changes in ambient temperature, although the speed of chemical reactions is temperature dependent (Sweeney and Hastings (1960)). This temperature compensation, with Q_10_ temperature coefficients of as little as 0.85 to 1.2, is a defining feature of circadian clocks (Pittendrigh (1960)) and has been proposed to be part of a general homeostatic mechanism that confers robustness to the circadian timing system under a wide range of unstable environmental conditions (Pittendrigh and Caldarola (1973); Sancar et al. (2012); Johnson and Egli (2014)). However, the molecular mechanisms underlying circadian temperature compensation are still poorly understood.

Using a large-scale RNAi screen, we identified cleavage and polyadenylation specificity factor 6 (*CPSF6*, also known as *CFIm68*) as a regulator of circadian temperature compensation. *CPSF6* is a core component of the cleavage and polyadenylation machinery required for proper posttranscriptional processing of mRNA molecules. Interestingly, posttranscriptional regulation in mediating temperature perception by circadian clocks has been reported in several organisms. In *Drosophila*, thermosensitive alternative splicing of the *period* gene plays a critical role in the adaption to seasonally cold days (Majercak et al. (1999)). Similarly, temperature-associated alternative splicing coupled to nonsense mediated decay regulates the expression of several core clock genes in *Arabidopsis thaliana* (James et al. (2012); Kwon et al. (2014)). In addition, temperature affects translational efficiency of certain plant genes such as bHLH transcription factor PIF7 through thermosensitive conformational changes within its mRNA 5’ UTR, thus influencing thermomorphogenesis (Chung et al. (2020)). Furthermore, posttranslational protein modifications such as phosphorylation or sumoylation have been shown to be important regulators of temperature compensation in plants, mammals and fungi (Mehra et al. (2009); Portolés and Más (2010); Zhou et al. (2015); Hansen et al. (2017)).

For our RNAi screen, we therefore focused on genes associated with posttranscriptional modification and RNA processing and identified *CPSF6* as the gene with the most significant period phenotype among all those tested. At a physiological temperature of 37°C, knockdown of *CPSF6* leads to a strong dose-dependent lengthening of the circadian period in human U-2 OS and mouse NIH3T3 cells and, most importantly, a tremendously impaired temperature compensation. Together with *NUDT21* (also known as *CPSF5* or *CFIm25*), which shows a similar phenotype upon knockdown, *CPSF6* forms a tetramer that binds to UGUA motifs of pre-mRNAs (Brown and Gilmartin (2003)), thereby promoting the utilization of distal polyadenylation sites of its targets (Martin et al. (2012)). We found that knockdown of *CPSF6* altered 3’ UTR length, mRNA stability and expression of specific clock genes such as *CLOCK*, *NR1D2* or *PER3*. However, these properties did not show a systematic temperature-dependency, indicating a more complex regulation of temperature decompensation in *CPSF6* knockdown cells. Therefore, we have globally searched for key regulators controlling circadian temperature compensation through the regulation of posttranscriptional processes by extending our analysis to the transcriptome and proteome.

At 37° C, knockdown of *CPSF6* resulted in global shortening of 3’ UTR length in approximately one-third of all detected transcripts, similar to results previously reported for human embryonic kidney cells (HEK-293; (Martin et al. (2012); Zhu et al. (2018))) and mouse myoblasts (C2C12; (Li et al. (2015))). In addition, knockdown of *CPSF6* resulted in global changes in transcript and protein expression, but these can only be attributed to changes in 3’ UTR length to a limited extent, similar to previous observation (Gruber et al. (2014)). How might such *CPSF6*-dependent effects influence circadian temperature compensation? Two conceptually different hypothesis have been proposed as the mechanism underlying temperature compensation. Whereas the critical reaction hypothesis assumes that period-determining reactions are insensitive to temperature, the balance hypothesis suggests a counterbalance of period lengthening and shortening effects upon temperature changes. Regardless of which hypothesis is correct, we postulated that the temperature response after knockdown of *CPSF6* would be reflected in differential temperature-dependent behavior of either 3’ UTR length, transcript or protein expression. Indeed, we identified several genes, whose temperature response is *CPSF6*-dependent at at least one of the three regulatory levels, but interestingly, only six genes that are regulated by temperature and *CPSF6* at all three levels (at our chosen significance thresholds). Among these six genes, eukaryotic translation initiation factor 2 subunit 1 (*EIF2S1*, also known as *eIF2α*) showed the most significant circadian period phenotype in our large-scale RNAi screen.

Interestingly, *EIF2S1* has been described to be a key sensor for a variety of stress conditions including temperature, metabolic fluctuations and other environmental challenges(Boye and Grallert (2020)). Cellular stress results in phosphorylation of EIF2S1 by four different EIF2S1 kinases, which integrate various stress signals into a common pathway often resulting in attenuation of global translation and activation of translation of some mRNAs such as *ATF4* (activating transcription factor 4). For example, low temperature activates the EIF2S1 kinase GCN2 (general control non-derepressible 2/eIF2αkinase 4) (Roobol et al. (2015)), whereas high temperature activates the EIFS1 kinase PERK (PKR-like eukaryotic initiation factor *2α* kinase) (Park et al. (2018)).

EIF2S1 does not only act as a master sensor of intrinsic and extrinsic stress signals, it also has been described to control circadian rhythms in diverse organisms. In mammals, EIF2S1 phosphorylation levels regulate the circadian period by promoting translation of *Atf4* mRNA, a transcription factor targeting binding motifs within the promoters of several clock genes (Pathak et al. (2019)). In addition, clock controlled rhythmic accumulation of phosphorylated EIF2S1 translationally controls rhythmic gene expression in *Neurospora crassa* (Karki et al. (2020)). In mouse liver, rhythmic *EIF2S1* expression mediates circadian regulation of stress granula (Wang et al. (2019)).

Recently, many transcripts have been shown to exhibit circadian oscillations in 3’ UTR length (Gendreau et al. (2018); Greenwell et al. (2020); Yang et al. (2020)). It would be interesting to find out in future studies whether and to what extent the rhythmicity in 3’ UTR length affects the mechanisms of temperature compensation. In our study, we showed that a small fraction of the transcriptome exhibits temperature-dependent changes in 3’ UTR length. Thus, the previously reported daily rhythms in alternative polyadenylation may be additionally regulated by temperature, potentially allowing the system to plastically shape the circadian transcriptome in response to changing environmental conditions. Our study suggests that alternative polyadenylation is a posttranscriptional mechanism underlying temperature compensation of circadian periods in human cells. Alternative polyadenylation has been shown to be highly tissue dependent in diverse organisms such as mammals (Gruber et al. (2012); Lianoglou et al. (2013); Floor and Doudna (2016)) or insects (Smibert et al. (2012)), very similar to properties of circadian systems (Menet and Hardin (2014); Endo et al. (2014); Pett et al. (2018)). It would therefore be interesting to investigate in future studies whether alternative polyadenylation acts as an additional regulatory layer to confer tissue-specific fine-tuning of circadian clocks.

Taken together, we here provide evidence for posttranscriptional control of circadian temperature compensation in mammals. We found a key regulator of polyadenylation site selection and hence 3’ UTR lengths, *CPSF6*, to modulate not only the period of the circadian oscillator but also the transcription and translation of a huge variety of genes in a temperature-dependent manner. This includes the master regulator of cellular stress response, *EIF2S1*, whose transcript and protein level depend on both *CPSF6* and temperature. *EIF2S1* is not only a temperature and metabolic sensor, but also a clock regulator making it a very plausible candidate for integrating and processing fluctuating environmental signals to establish a robust circadian period and phase. Future studies are needed to characterize these properties in detail.

## Materials and Methods

### RNAi screen

Large scale RNAi-based screening for circadian phenotypes has been done as previously described (Maier et al. (2009)). In essence, RNAi-constructs have been purchased from Open Biosystems, Lentiviruses were produced in HEK293T cells in a 96-well plate format as described earlier (Brown et al. (2008)) and U-2 OS cells harboring a *Bmal1:Luc* reporter construct were transduced with a 100μl virus filtrate plus 8ng/μl protamine sulfate. Oscillatory properties of bioluminsescence recordings have been estimated using the ChonoStar software, which detrends the raw data by a 24h-window moving average filter and subsequently fits the detrended data to an exponentially damped cosine function (Maier et al. (2021)). Statistical properties of circadian oscillatory parameters such as amplitudes, damping coefficients and periods based on multiple RNAi constructs per gene are assessed via a Redundant SiRNA Activity (RSA) analysis as previously described (König et al. (2007)).

### Sample Collection

U-2 OS BLH cells were transduced with RNAi lentiviruses with bulk method. In brief, U-2 OS cells grown in ten T175 flasks were taken in suspension by trypsin digest and evenly distributed to eight 50 mL Falcon tubes. Pelleted cells were carefully resuspended in 4.5 mL lentiviral supernatant (either pGIPZ non-silencing or pGIPZ targeting *CPSF6*, V3LHS_640891) containing protamine sulfate (at 8μg/mL final concentration). Cells are spun at 37°C, 800 x g for 90 minutes. After centrifugation, cells are gently resuspended using a 1 mL pipette and seeded to 3 × T75 flasks per 50 mL Falcon tube (in total 24 × T75 flasks) and incubated at 37°C. 5d after transduction, cells were transfered to respecitve temperature conditions (32°C, 37°C or 39°C). Cells were harvested for RNA or protein isolation 4 days after transfer to respective temperature conditions.

### Protein extraction

Subsequent procedures have been performed on ice or at 4°C. Cells grown in cell culture flasks (75cm^2^) were lysed by adding 0.5ml of lysis buffer containing 4% SDS in 100mM Tris pH 8.5. Lysates were collected with cell scraper, transfered to Eppendorft tubes, boiled for 15 min at 95^°^C and stored at −80°C.

### RN A-sequencing

Two protocols have been applied to pursue different goals: To precisely map polyadenylation sites in wild type U-2 OS cells we applied the 3’Seq RNAseH^+^ protocol as previously described (Afik et al. (2017)). To quantify overall transcript expression and 3’ UTR length (NCOM) in wild type and *CPSF6* knockdown cells we applied 3’end seq without RNAseH digest (RNAseH^−^) to preserve relative quantities of RNA fragments.

A schematic drawing of the analysis pipeline can be found in Fig. S10

#### RNAseH^+^ protocol

U-2 OS wild type and *CPSF6* knockdown cells are cultured at 37°C. After 4 days, cells are harvested and RNA extracted using a standard isolation kit (Invitrogen, RNeasy), Zn-fragmented and fragments with poly(A) tails are selected using oligo-(dT) beads. These pre-selected fragments are subsequently digested by an RNAse H treatment (RNAseH^+^-seq) which removes the poly(A)-RNA and allows for an exact linker ligation at the cleavage and polyadenylation site (Derr et al. (2016); Afik et al. (2017)). Strand specific reads of this 3’ end sequence library are aligned to the reference genome GRCh37 (hg19) and expression is quantified by the End Sequence Analysis Toolkit (ESAT, (Derr et al. (2016))). Polyadenylation signals are searched up to 5000bp beyond the annotated end of transcripts.

#### RNAseH^−^ protocol

In case of newly identified polyadenylation sites through our RNAseH^+^ protocol we extended the reference genome annotation by the newly identified polyadenylation site. Extended isoforms are markedby ’_ext’ to the RefSeqID in Tab. S3, S4, S6, S7, S9, S10. This new annotation has then been used for a further 3’ UTR expression analysis. 3’end identification and expression quantification has been again done via ESAT, using a scanning window length of 50 bases. Reads are subsequently normalized using DESeq (Anders and Huber (2010)) and transcript isoforms with identical 3’ UTRs are merged together.

### Proteomics

Four independent replicates of U-2 OS cell cultures at 3 temperature conditions for wild type and *CPSF6* knockdown cells were subjected to quantitative MS-based proteomics analysis. In brief, frozen cell lysates in 4% SDS buffer were prepared for MS-based quantitative proteomics analysis. Briefly, samples were boiled for 5 minutes at 95°C and homogenates were sonicated using a Bioruptor (2 × 15 cycles at 4°C; 30sec ON, 30sec OFF each cycle, at maximum power). Once a homogeneous solution was formed, the protein concentration was estimated based on Tryptophan assay. 1mg of protein from each sample was used as starting material for protein digestion. For this, protein lysates were incubated for 20 minutes at RT with 2.5mM Dithiothreitol (DTT) and 27.5mM 2-Chloroacetamide (CAA) for protein reduction and alkylation, respectively. The lysates were then precipitated with 80% acetone (overnight incubation at −20°C) and collected the following day by centrifugation (1,500 xg, 10 minutes, 4°C). After the protein pellets were washed four times with ice-cold 80% acetone, and air-dried at RT for 20 minutes, 500μL of trifluoroethanol (TFE) solution was added to each sample. Pellets were resuspended by sonication using a Bioruptor (15 cycles at 4°C 30sec ON, 30sec OFF each cycle, at maximum power), and subsequently incubated with endopeptidase LysC for 1h at RT followed by overnight incubation with Trypsin at 37°C. For both digesting enzymes the proportion to the protein amount was 1:100. After digestion, the samples were centrifuged (10,000 xg, 10 minutes, RT) to remove potential cellular debris, and 20μg were used for peptide desalting and clean-up using SDB-RPS StageTips. After eluting (80% ACN, 2.5% NH4OH), the peptides were concentrated in a centrifugal evaporator (45°C) until dryness and were reconstituted in LC-MS loading buffer (2% ACN, 0.2% TFA) and the eluate was stored at −20°C. A volume corresponding to 400ng of peptides was used for the analysis using an LC 1200 ultra-high-pressure system (Thermo Fisher Scientific) coupled via a nano-electrospray ion source (Thermo Fisher Scientific) to a Q Exactive HF-X Orbitrap (Thermo Fisher Scientific). Prior to MS, the peptides were separated on a 50cm reversed-phase column (diameter of 75mm packed in-house with ReproSil-Pur C18-AQ 1.9mm resin [Dr. Maisch GmbH]) over a 120min gradient of 5%-60% buffer B (0.1% formic acid and 80% ACN). Full MS scans were acquired in the 3001,650m/z range (R=60,000 at 200m/z) at a target of 3e6 ions. The fifteen most intense ions were isolated, fragmented with higher-energy collisional dissociation (HCD) (target 1e5 ions, maximum injection time 120ms, isolation window 1.4m/z, NCE 27%, and underfill ratio of 20%), and finally detected in the Orbitrap (R=15,000 at 200m/z). Raw MS data was processed with MaxQuant and reported LFQ normalized protein intensities used for further bioinformatic analysis.

## Acknowledgments

We gratefully acknowledge Astrid Grudziecki and Maike Mette Thaben for excellent technical assistance as well as Paul Thaben for support in statistical analysis.

## Funding

CS acknowledges support by the Deutsche Forschungsgemeinschaft (DFG, German Research Foundation) through grant number SCHM3362/2-1. SK acknowledges support by the National Institute of Health under award number R01GM125859. Work in AK’s and HH’s laboratory is funded by the Deutsche Forschungsgemeinschaft (DFG, German Research Foundation) - Project-ID 278001972 - TRR 186. We thank the LMU Munich’s Institutional Strategy LMUexcellent within the framework of the German Excellence Initiative and the Deutsche Forschungsgemeinschaft (DFG, German Research Foundation) SFB1321 (Project-ID 329628492) for funding the work of MSR.

## Conflict of Interest

The authors declare that they have no conflict of interest.

## Notes

### Competing Interest Statement

The authors have declared no competing interest.

