## Supplementary Material for "An integrative omics approach reveals posttranscriptional mechanisms underlying circadian temperature compensation"

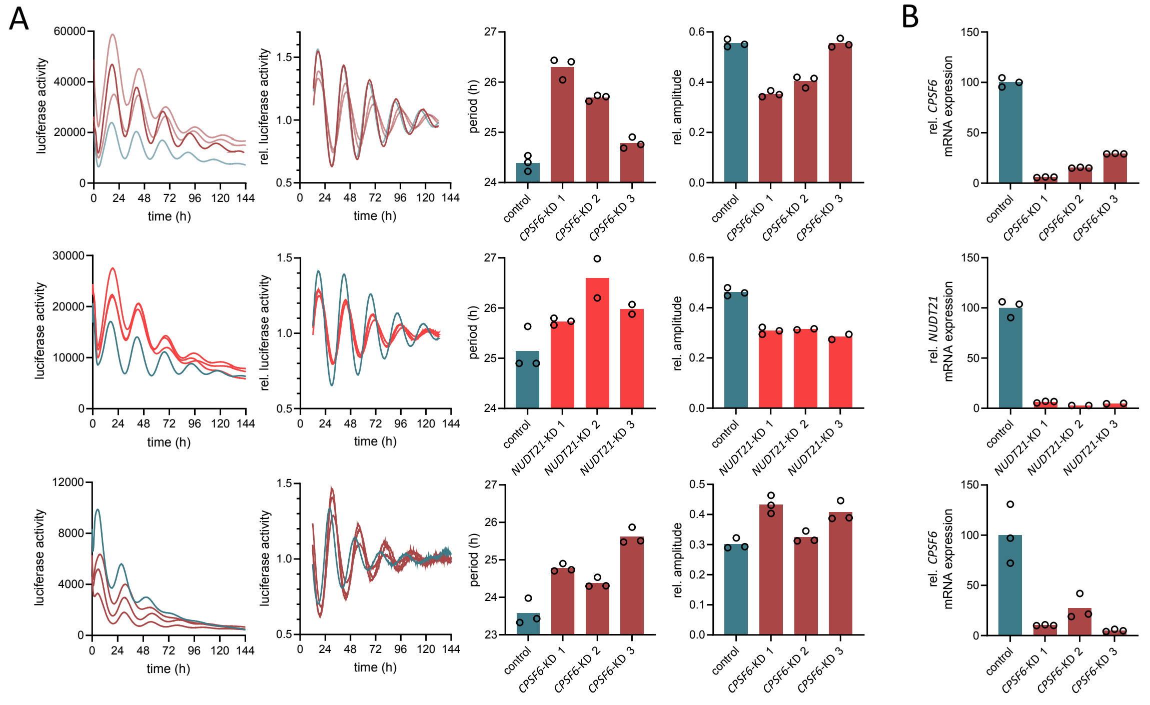


**Figure S1: Circadian free running period dose-dependently lengthens upon *CPSF6* knockdown.** *Upper row*: A) Raw (1st column) and detrended (2nd bolumn) *Bmal1-luciferase* expression in human wild type (*blue*) and *CPSF6* knockdown (*red*) U-2 OS cells, using three different siRNA constructs. Periods (3rd column) and relative amplitudes (4th column) dose-dependently (B) lengthen or decrease, respectively, upon *CPSF6* knockdown. *Middle row:* Analogous behavior can be observed for *NUDT21* (*red*), a heterodimeric binding partner of *CPSF6*. *Bottom row:* A dose-dependent lengthening of period and increase of relative amplitude of *Bmal1-luciferase* expression upon *CPSF6* knockdown can be observed in mouse NIH3T3 cells, analogous to findings in human U-2 OS cells (*upper panel*).

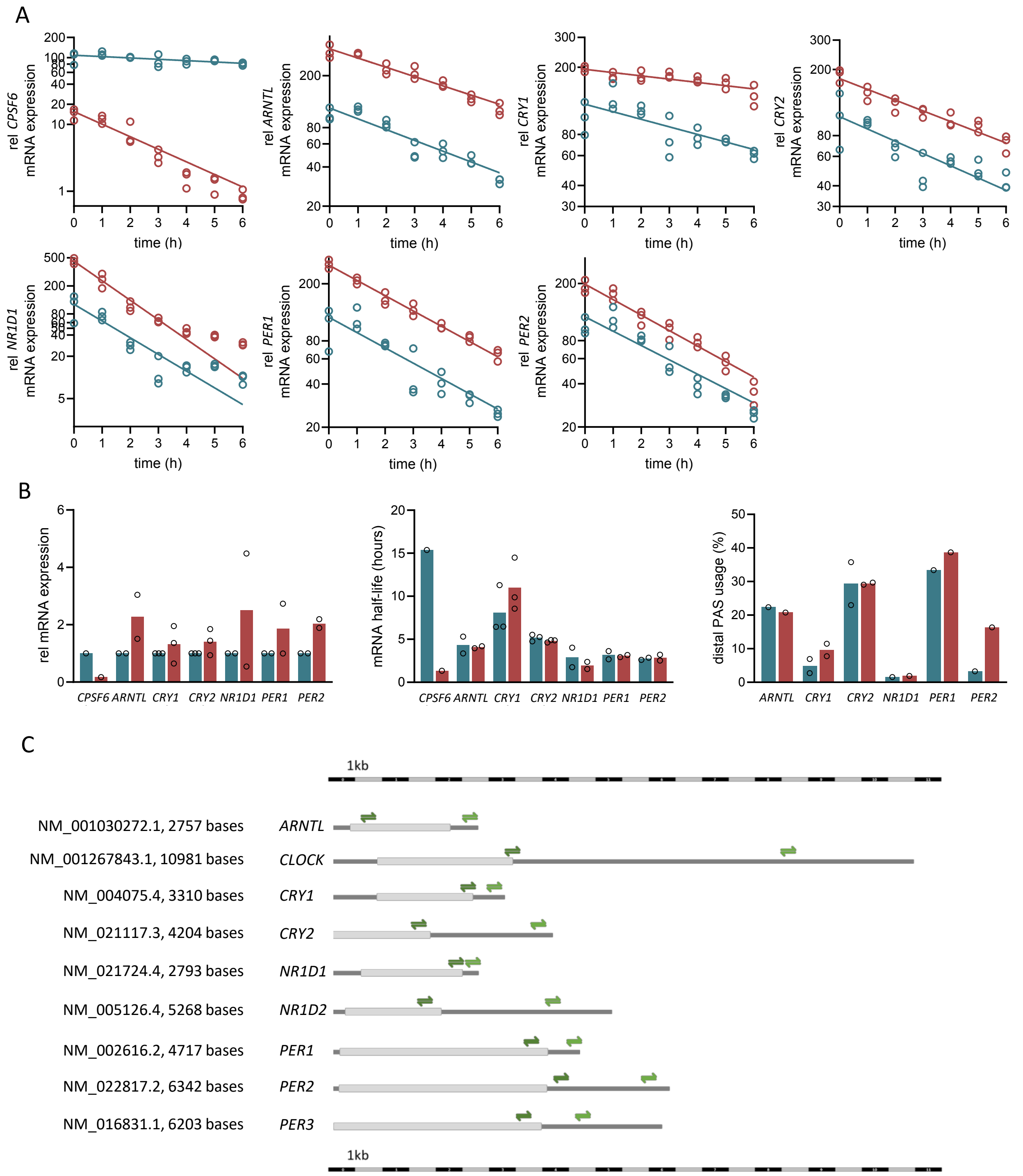


**Figure S2: *CPSF6* knockdown induced changes in poyladenylation site usage, mRNA expression and mRNA half-life among canonical clock genes.** A-B) Analogous to Fig. 1B *middle panel* and *bottom panel* of the *Main text*, respectively, showing results for *CPSF6*, *Arntl*, *Cry1*, *Cry2*, *Nr1d1*, *Per1* and *Per2*. C) Schematic representation of primer location, used to measure expression of short- and long-3' UTR isoform expression of nine core clock genes.

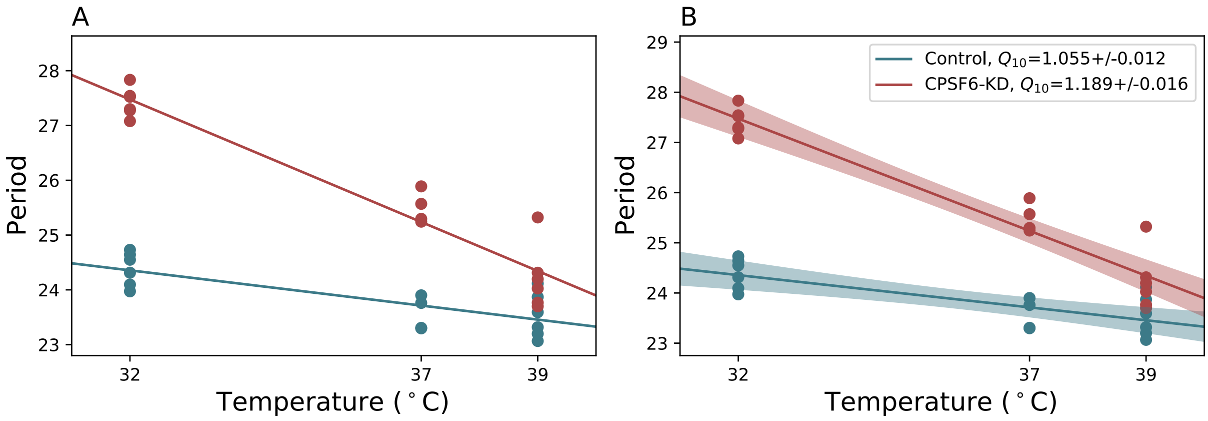


**Figure S3: Statistical accession of differential temperature response and calculation of period temperature coefficient $Q_{10}^{\tau}$.** A) Temperature response is statistically different in wild type and *CPSF6* knockdown cells, assuming a linear dependency of free-running period $\tau$ on temperature $T$. The test for statistically significantly different slopes in two groups of data has been done as outlined in Chapter 11.4 of (Armitage et al. (2002)). B) Calculation of $Q_{10}$ temperature coefficients from slopes, determined in panel (A). Temperature coefficients have been obtained, using the equation $Q_{10} = \left(\frac{\tau_1}{\tau_2}\right)^{10^\circ\mathrm{C}/7^\circ\mathrm{C}}$ with $\tau_1$ and $\tau_2$ being the free-running period determined at 32°C and 39°C as obtained from the linear regression, respectively. Error propagation has been calculated via the Python uncertainties package.

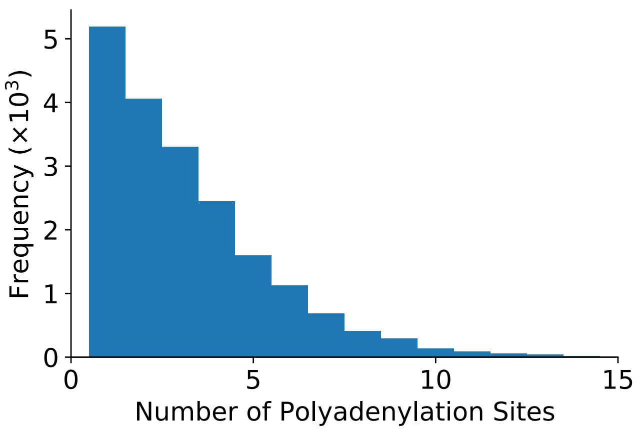


**Figure S4: Number of polyadenylation sites per gene as detected by the RNAseH$^-$ protocol.**

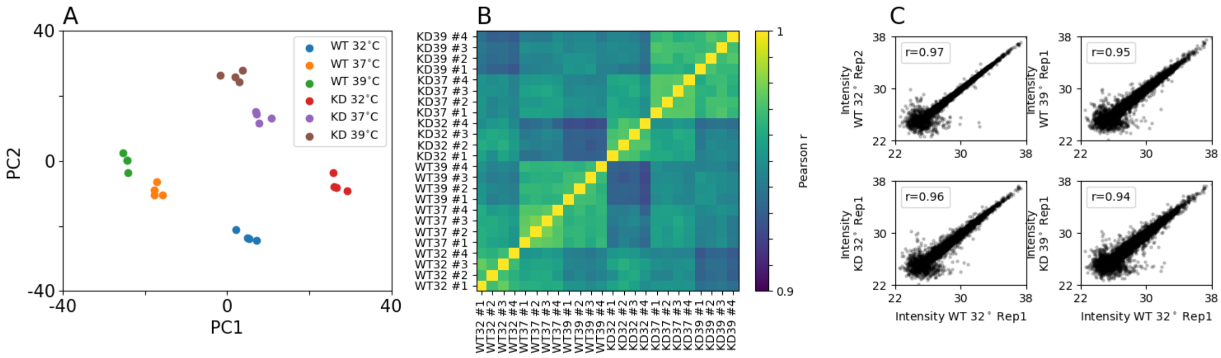


**Figure S5: Quality control of proteomics data.** A) Principal component analysis on log2 transformed protein LFQ intensities as determined by the MaxQuant software (Cox and Mann (2008)). Technical replicates as indicated by equal coloring group together. B) Correlogram of the Pearson correlation coefficients (r) from the log2 transformed protein LFQ intensities across measured samples. As expected, replicates with the same genetic background (wild type versus *CPSF6* knockdown) and at the same environmental temperature show the highest correlations of abundance values. C) Representative scatter plots of log2 transformed protein LFQ intensities across biological replicates with the Pearson correlation coefficient ($r$).

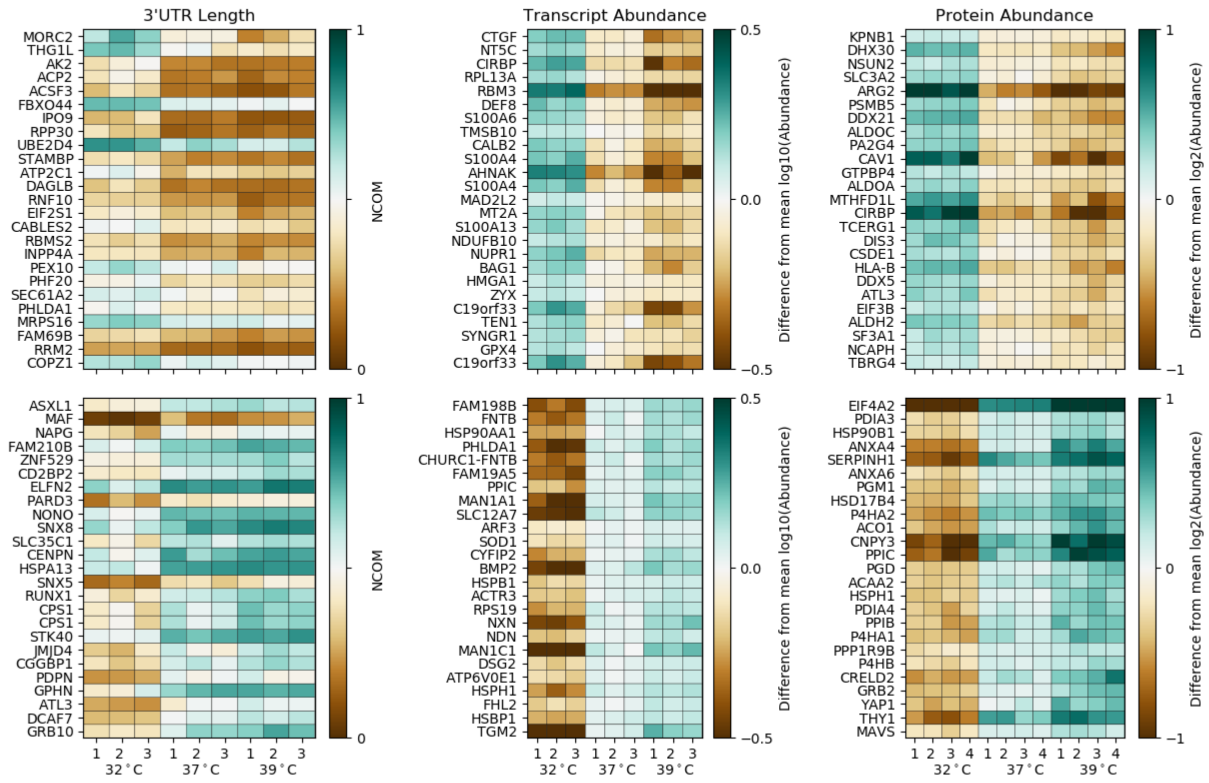


**Figure S6: Genes with most significant variations in 3' UTR length, transcript expression and protein abundance upon changes in environmental temperature.** A) Heatmap of 25 NCOM values that exhibit the most significant shortening (*upper* panel) or lengthening (*bottom* panel) of 3' UTRs upon changes in environmental temperature in *wild type* U-2 OS cells, sorted by increasing Benjamini-Hochberg corrected p-values. All replicates at all three temperatures, namely 32°C, 37°C and 39°C, are shown. B) Same as panel (A), showing the 25 genes with the most significant down- (*upper* panel) or up-regulation (bottom) in gene expression upon increasing environmental temperatures. C) Same as panel (A) and (B), showing the 25 genes with the most significant down- (*upper* panel) or up-regulation (bottom) in protein abundance upon increasing environmental temperatures.

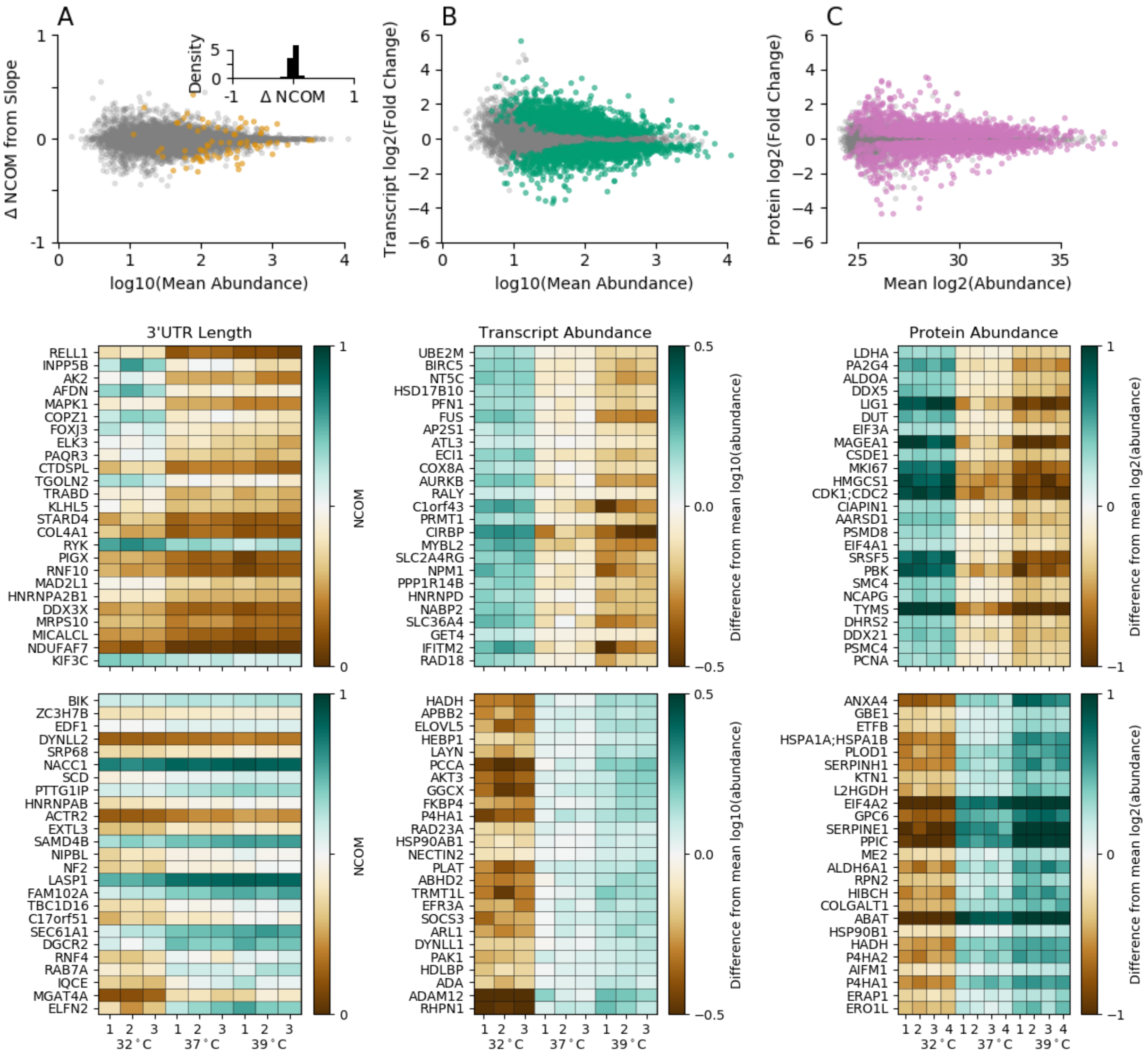


**Figure S7: Changes in 3' UTR length, transcript abundance and protein abundance upon temperature changes in *CPSF6* knockdown cells.** A-C) Same as Fig. 4A-C in case of *CPSF6* knockdown cells. D-F) Same as S6A-C in case of *CPSF6* knockdown cells.

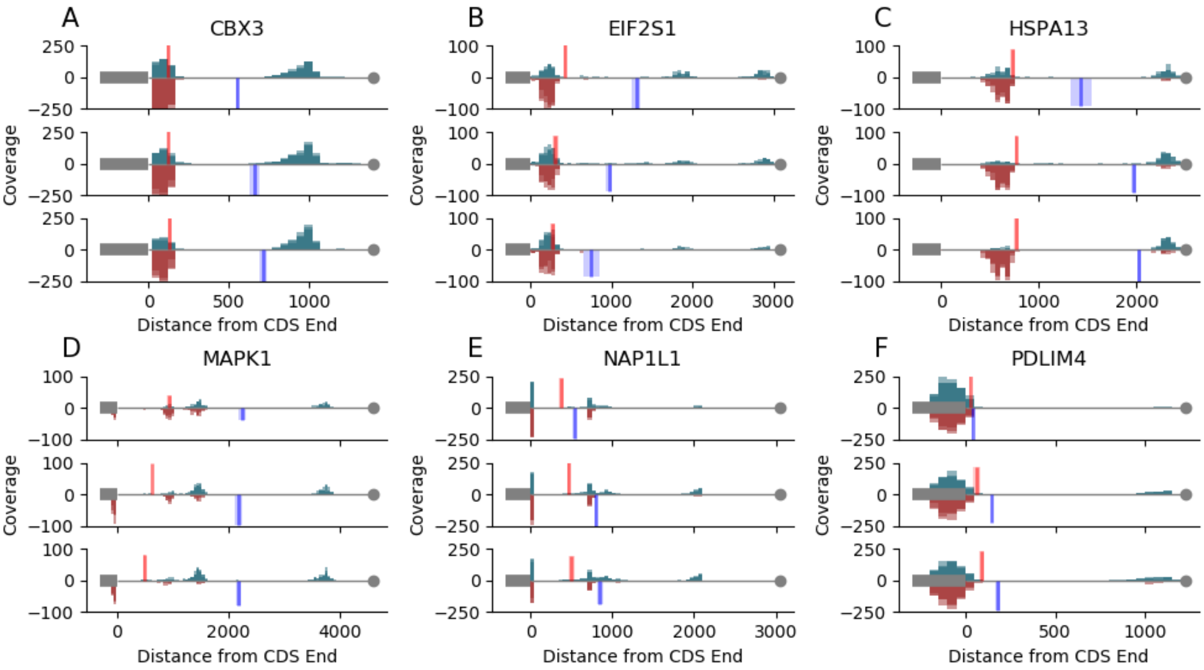


**Figure S8: Read distribution for genes exhibiting differential temperature responses between wild type and *CPSF6* knockdown cells.**

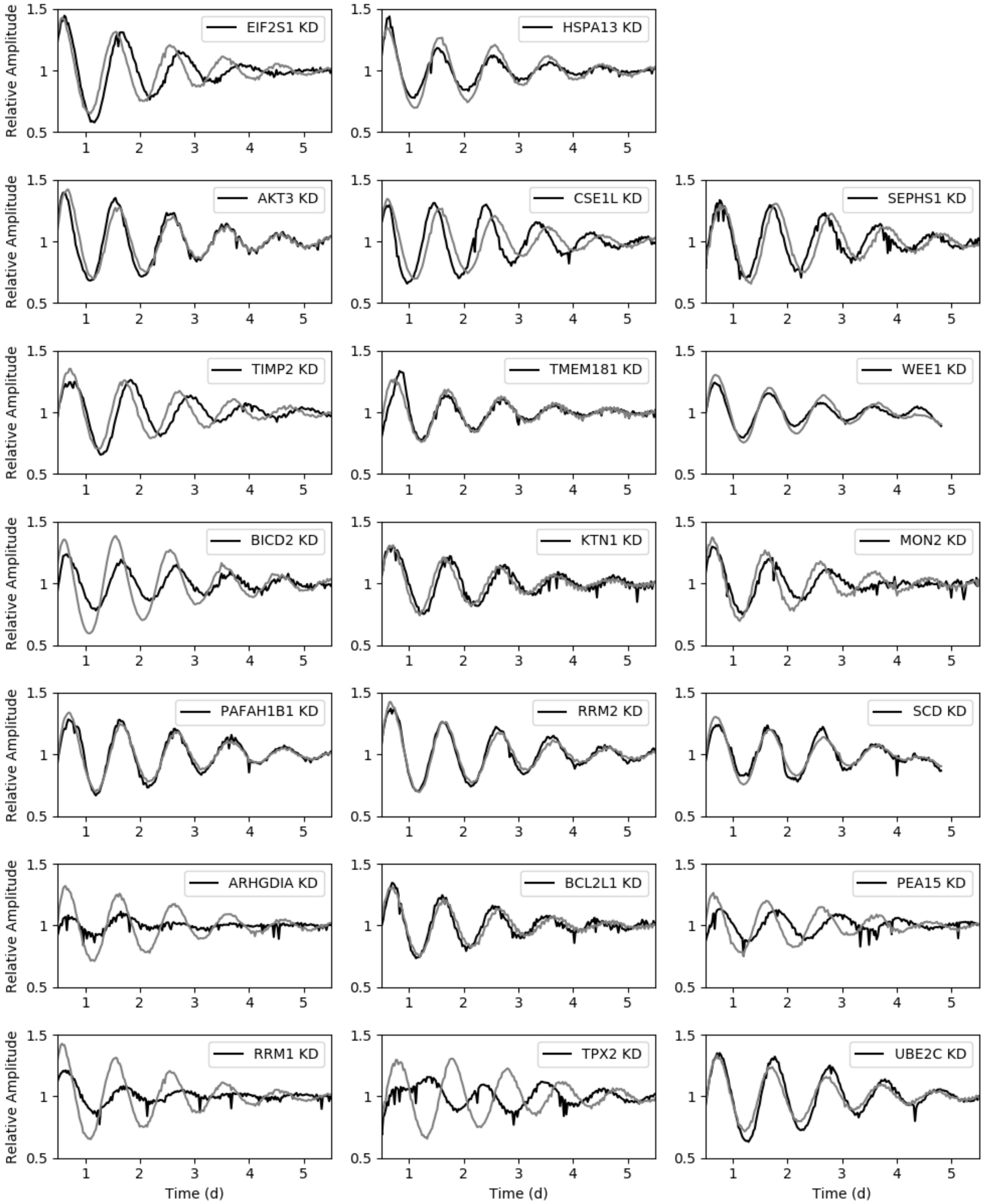


**Figure S9: Large scale RNAi screen example time traces.** Relative amplitude of *Bmal1-luciferase* oscillations for RNAi constructs (*black*) and the corresponding control (plate mean; *gray*). Depicted are examples for genes showing a significant period shortening or lengthening in the RNAi screen as well as a differential temperature response at the NCOM, transript and protein level (first row), the NCOM and transcript level (second and third row), the NCOM and protein level (fourth and fifth row) or the transcript and protein level (sixth and seventh row), compare Fig. 5D.

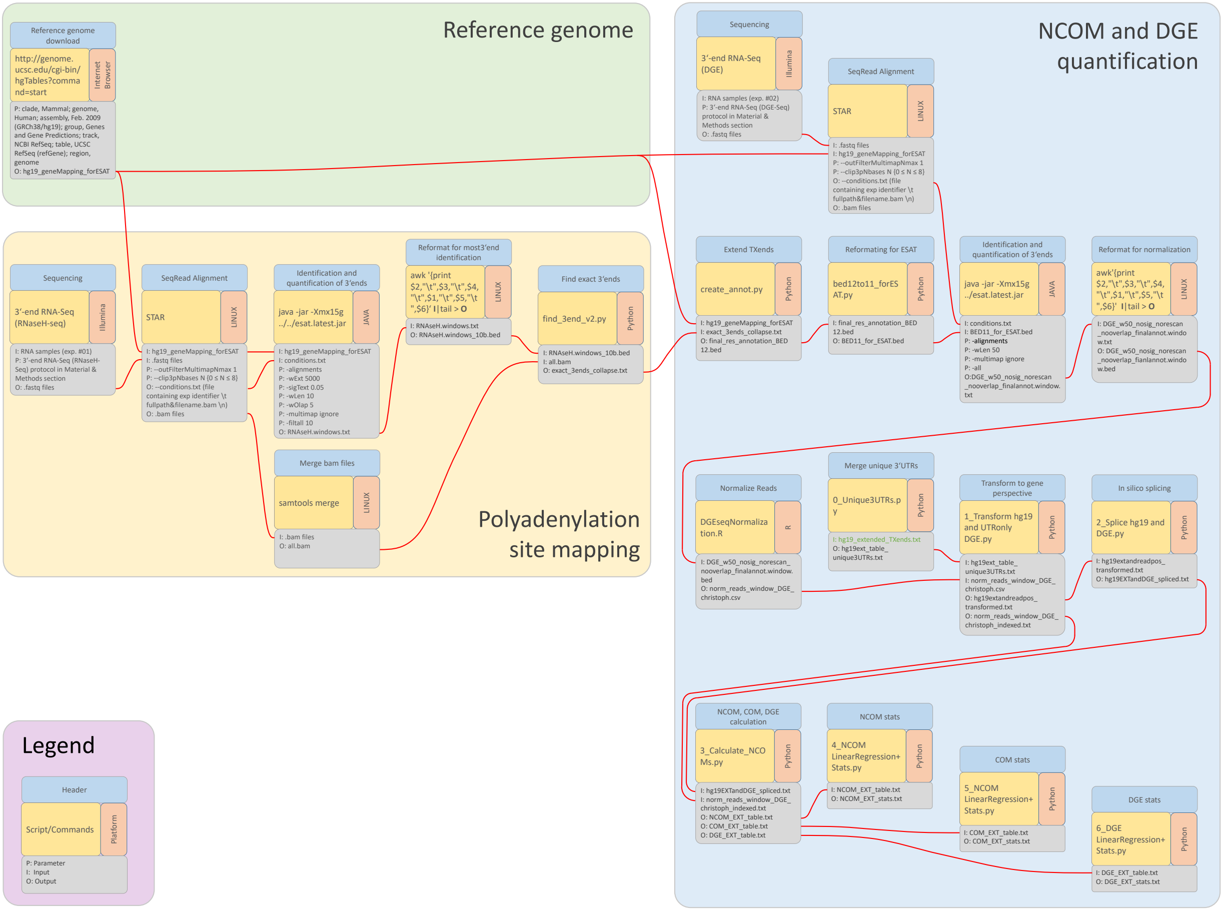


**Figure S10: Schematic drawing of the transcriptomics analysis pipeline.**

**Table S1: Oscillatory properties of *Bmal1-luciferase* expression within the large scale RNAi screen.**

**Table S2: Detected polyadenylation sites by the RNAseH-seq approach.**

**Table S3: Isoforms showing a significant shift in 3' UTR length (NCOM) upon *CPSF6* knockdown at 37°C.**

**Table S4: Isoforms showing a significant differential transcript expression between wild type and *CPSF6* knockdown cells at 37°C.**

**Table S5: Proteins showing a significantly altered abundance in wild type and *CPSF6* knockdown cells at 37°C.**

**Table S6: Isoforms showing a significant shift in 3' UTR length upon temperature changes in wild type cells.**

**Table S7: Isoforms showing a significant alteration in transcript expression upon temperature changes in wild type cells.**

**Table S8: Proteins showing a significantly altered abundance upon temperature changes in wild type cells.**

**Table S9: Isoforms showing a significantly differential response of 3' UTR length between wild type and *CPSF6* knockdown cells upon temperature changes.**

**Table S10: Isoforms showing a significantly differential response of transcript expression between wild type and *CPSF6* knockdown cells upon temperature changes.**

**Table S11: Proteins showing a significantly differential response of abundance between wild type and *CPSF6* knockdown cells upon temperature changes.**
